# Interactions between the visual and motor systems during optokinetic stimulation and simultaneous dynamic handgrip exercise: an EEG study

**DOI:** 10.1101/2020.03.31.011734

**Authors:** Fernando Cross-Villasana, James Dowsett, Angela Mastropasqua, Marianne Dieterich, Paul C.J. Taylor

## Abstract

Inhibitory interactions between the visual and motor systems are robustly found during the transient stimulation of either system. However, it is unknown how such interactions operate if both systems are continuously and simultaneously activated. To test this, we engaged the visual-oculomotor system in 14 right-handers with a large moving striped pattern that elicits characteristic eye movement responses, the optokinetic nystagmus (OKN). We also engaged the motor system with dynamic handgrip exercise simultaneous to OKN. We hypothesised that central versus occipital EEG upper-alpha (10-12 Hz) synchronisation would reflect systems-level inhibition between visual and motor processes. Optokinetic responses were recorded with eye tracking, and gripping force measured with dynamometry. First, when exploring whether continuous visual activation inhibits motor activity, we found that OKN increased alpha synchronisation over central electrodes, in line with classic findings of motor downregulation to transient visual stimuli. We found further evidence for the visual inhibition of motor-cortical activity, by comparing between visual conditions: increasing visual input, from closed eyes, to eye opening, to visual stimulation, produced progressively greater motor synchronisation into the inhibitory upper-alpha band. Second, we tested for the opposite interaction, that is, effects of motor engagement over ongoing visual and oculomotor processes. Behaviourally, left hand-grip exercise accelerated the ongoing OKN, increasing slow phase velocity. Electrophysiologically, performing this repetitive motor task with either hand interfered the occipital alpha desynchronization that OKN normally induces on EEG. These results support that mutual visual and motor inhibitory interactions persist during their simultaneous engagement, and show their effects on eye movements.

**Key points:**

- Increasing levels of visual activation produced increasingly stronger signs of cortical inhibition.
- Hand motor activation interfered with visual cortical activation during visual stimulation.
- Hand motor activation altered oculomotor responses.
- Ongoing visual-motor interactions were reflected by the EEG alpha band.
- EEG can be reliably assessed during nystagmus.

## 1. Introduction

Co-ordinating between all the possible tasks that we are capable of at any given moment requires a large-scale system for striking the right balance of facilitation and downregulation on the involved processes in the brain. Inhibitory interactions between the visual and the motor systems have been classically shown when either one of them is engaged transiently and /or in isolation (e.g. Becker-Bense et al., 2006, Cross-Villasana et al., 2016, Delorme et al., 2007, Koshino and Niedermeyer, 1975, Pfurtscheller and Lopes da Silva, 1999). The function of these interactions is presumed to facilitate attention to the stimulated system by down-regulating the other system (Crabtree and Isaac, 2002, Kober et al., 2015, Sterman, 1996). This leaves the open question as to how these interactions play out during ongoing behaviour (Huk et al., 2018) where both, the visual and the motor systems are engaged continuously and simultaneously.

As seen in classic EEG studies, the execution of isolated hand movements leads not only to activation of the motor cortex but also simultaneous down-regulation of the visual cortex (e.g. Gerloff et al., 1998, Pfurtscheller and Lopes da Silva, 1999). This phenomenon can also be observed over longer time periods during the execution of dynamic handgrip exercise (Cross-Villasana et al., 2016) or during sequential button-pressing (Deiber et al., 2001). The opposite phenomenon, where visual processes down-regulate motor activity, has been observed in the EEG when a word to read is presented onscreen (Pfurtscheller and Lopes da Silva, 1999), during visual pattern exploration (Koshino and Niedermeyer, 1975) or through the presentation of flickering lights (Brechet and Lecasble, 1965). Brain stimulation studies have suggested that increasing visual activation even just by opening the eyes reduces motor excitability (Leon-Sarmiento et al., 2005, Strigaro et al., 2015).

Electrophysiological evidence suggests that mutual inhibitory interactions arise at the thalamic level, where different sensory systems can exert inhibitory modulation on each other via the thalamic reticular nucleus (TRN; Crabtree and Isaac, 2002, Edlow et al., 2012, Guillery and Harting, 2003, Zikopoulos and Barbas, 2007). In this account, when a thalamic nucleus relays a signal towards the cortex, its axon collaterals simultaneously stimulate sections of the TRN that in turn down-regulate relay nuclei from other systems (Crabtree and Isaac, 2002, Guillery and Harting, 2003, Zikopoulos and Barbas, 2007). As the TRN also mediates the generation of the cortical alpha rhythm (Hughes and Crunelli, 2005, Larson et al., 1998, Vijayan and Kopell, 2012), the result of inter-sensory inhibition may be reflected in the scalp EEG as an alpha power enhancement over the inhibited cortex (Pfurtscheller and Lopes da Silva, 1999, Steriade, 2000, Sterman, 1996). Furthermore, it has been observed that stimulation of one thalamocortical pathway can interrupt ongoing activity in another pathway through TRN-mediated inhibition in the thalamus (Crabtree and Isaac, 2002, Zikopoulos and Barbas, 2007). Therefore, such interference between concurrently-active thalamocortical streams should also be observable over scalp EEG alpha measurements. This proposal is here examined by testing for the classic visual down-regulation produced by hand-movements, but during active visual stimulation. Modulation of the visual and motor systems throughout can then be tracked by time-frequency analysis of the EEG alpha band. In particular, in the upper-alpha band, amplitude enhancements are indicative of cortical inhibition and amplitude decrements reflect activation (Hummel et al., 2002, Klimesch et al., 2007, Pfurtscheller and Lopes da Silva, 1999, Zarkowski et al., 2006).

Here we implement steady activation of the visual system through optokinetic stimulation (OKS), which is then paired with simultaneous motor activation through dynamic-handgrip exercise (Kluess et al., 2000) to interfere with that visual activation (Cross-Villasana et al., 2016). By itself, OKS has reliably induced visual-cortical activation in EEG (Gulyas et al., 2007, Magnusson et al., 1985) and fMRI studies (Becker-Bense et al., 2006, Bense et al., 2001, Dieterich et al., 2003, Kikuchi et al., 2009, Ruehl et al. 2019). In the current study, OKS consists of a series of moving stripes presented on a screen, which induce a reflexive eye movement that repetitively tracks their motion (Ventre-Dominey and Luyat, 2008), called optokinetic nystagmus (OKN; Ilg, 1997). During OKN, the eyes follow the direction of OKS through a slow-phase movement, which is followed by a quick reset in the opposite direction, called the quick-phase, followed again by a slow phase, and so on (Ilg, 1997, Ventre-Dominey and Luyat, 2008).

The OKN itself may be affected by dynamic handgrip, though different predictions can be made according to diverging accounts. On the one hand, given the downregulation of the visual cortex that dynamic handgrip have previously been found to produce (e.g. Cross-Villasana et al., 2016), this could theoretically disrupt the engagement of the subsequent oculomotor mechanisms that produce nystagmus, leading to a reduction of the OKN slow-phase velocity. On the other hand, stimulation of other sensory modalities, such as acoustic or tactile, has been shown to increase the OKN slow-phase velocity (Magnusson et al., 1985). This could be the case with dynamic handgrip, which involves tactile input.

Next we turn to the prediction of how OKS may affect the motor system. In an OKS-fMRI study (Becker-Bense et al. (2006), it was observed that apart from visual activation, OKS also down-regulated motor areas, similar to the effects seen in EEG studies with different visual stimuli (Brechet and Lecasble, 1965, Koshino and Niedermeyer, 1975, Pfurtscheller and Lopes da Silva, 1999). However, previous OKS-EEG studies did not assess anterior regions (Gulyas et al., 2007, Magnusson et al., 1985) hence it is unclear whether any inhibitory processes (alpha power increases) are observable at central and anterior locations during OKS. This absence of evidence may also be due to the problem of the unavoidable eye-movement artefacts in the EEG induced by OKN (Gulyás et al., 2007), which are stronger at anterior regions. However, recent advances allow circumventing this problem through techniques such as independent component analysis (ICA), used to separate out eyemovement components (Bell and Sejnowski, 1995, Makeig et al., 1995) and speculated to be potentially as applicable to OKN EEG artefacts as to saccades (Einhäuser et al. 2017). Spatial filters like the surface Laplacian further reduce volume conduction and help to specify the location of experimental effects (Kayser and Tenke, 2015, Tenke and Kayser, 2005). Therefore, to deal with eye movements, those techniques are extended in the current design, building on a previous study (Cross-Villasana et al., 2018). Within the current design, detecting the classic bilateral motor activation (alpha reduction) produced by hand movements (e.g. Cross-Villasana et al., 2016, Pfurtscheller and Lopes da Silva, 1999) but during simultaneous OKN, would support the reliability of EEG measurement. With EEG reliability demonstrated, we then proceed to investigate our main experimental aims: the effect of motor activity on visual processing during OKS, its effects on OKN, and the inhibitory influence of isolated OKS over motor regions.

## 2. Materials and methods

### 2.1 Study design

We compared the effects of simple OKS exposure with the effects of simultaneous OKS/dynamic handgrip, on the EEG upper-alpha band (10-12 Hz), and on the slow-phase velocity of OKN. EEG oscillations were tracked throughout OKS using time-frequency analysis. OKN was assessed via eye-tracking. Electro-oculography (EOG), electromyography (EMG) and grip-force measurements were used as control variables. Our main experimental hypotheses are summarized in Table 1.

**Table 1.** Summary of the experimental hypotheses.

|  |  |
| --- | --- |
| *During simultaneous OKS-Dynamic Handgrip* |  |
| 1. | The motor system shows reduced upper-alpha amplitude (activates) during OKS + left or right Handgrip compared to OKS alone (supporting reliable EEG measurement during OKN). |
| 2. | The visual system shows increased upper-alpha amplitude (deactivates) during OKS + left or right Handgrip compared to OKS alone. |
| 3. | OKN slow-phase velocity is affected during OKS + left or right handgrip compared to OKS alone. |
| *During increasing visual input towards OKS alone* |  |
| 4. | The visual system shows reduced upper-alpha amplitude (activates) with increasing visual input, from eyes closed baseline, to eyes open baseline, to OKS. |
| 5. | The motor system shows increased upper-alpha amplitude (deactivates) with increasing visual input, from eyes closed baseline to eyes open baseline, to OKS. |

The experiment was designed to be short-lasting to prevent effects of tiredness or boredom on the measurements, especially on the EEG. It consisted of six blocks and two baselines with small breaks in between, lasting around 15 min altogether. Only right-handed participants were included in the study. Laterality was assessed with the Flinders Handedness Survey (Nicholls et al., 2013), where scores between 5 and 10 reflect right-handedness.

### 2.2 Participants

From nineteen recruited participants, five were excluded: one participant could not perform dynamic handgrip, one showed no clear nystagmus, one had an incomplete eyetracking recording, and two had incomplete EMG recordings. The final sample consisted of 14 participants (7 female) with a mean age of 26 years (range: 23-30 years). Their mean laterality quotient was 9.5 (range: 5-10) in the Flinders Laterality Inventory. The study was approved by the ethics committee at the medical faculty of LMU Munich.

### 2.3 Procedure

Before the experiment, two EEG resting baselines were measured: an eyes-closed baseline (BLC) and then an eyes-open baseline (BLO), each 60 secs long and performed while fixating a white cross. Afterwards, EMG and grip-force baselines were assessed, where participants performed a single grip of maximum voluntary contraction (MVC) with either hand on the digital dynamometer (Kluess, Wood, and Welsch 2000; Liu et al. 2003). The main experiment consisted of six blocks. In each block, first a white fixation cross appeared for 5 seconds at the center of the screen, followed by 45 seconds of OKS, followed again by 60 seconds of fixation cross presentation to dissipate possible task-aftereffects before the next block. A chin-rest was used to stabilize the participants’ head with their eyes at the center-level of the screen. Screen edges were covered with a black rectangular cardboard frame, leaving a 28° x 13° (visual angle) visible screen background. OKS was presented on an LCD monitor (1680 x 1050 pixel resolution) with a refresh rate of 60 Hz, placed at 40 cm from the participant. OKS stimuli consisted of 19 black (RGB: 0, 0, 0) and 19 white (RGB: 255, 255, 255) bars (bar width 2.3 cm/3.3°) which travelled rightwards at 33°/s. After each block, participants took a brief break until they felt ready for the next block. The first experimental block always consisted of OKS without handgrip. This was done to assure proper EEG characterization of modulations induced by OKS, without possible effects of tiredness, boredom, or aftereffects from previous blocks. The second and third blocks consisted of combined OKS with Left handgrip (OKS+L) and OKS with Right hand grip (OKS+R), order counterbalanced. After the third block, the process was repeated, with the fourth block always being OKS alone, and the fifth and sixth blocks of counterbalanced OKS+L and OKS+R. In the handgrip blocks, during the initial 5 seconds fixation cross, participants held the dynamometer passively with one hand, the other resting on their lap, palm facing down. Participants were then instructed to repeatedly squeeze the hand-dynamometer at OKS onset, firmly as if “trying to make a fist”, at approximately 2 Hz, with a force that they could sustain for 45 s. No displays of force and frequency of the handgrip were presented to the participant, since these might distract attention from the OKS and interrupt the ongoing OKN. The assessed grip-force and EMG data were analysed to ensure compliance with the task and to verify comparable effort exertion with each hand. During the last 60 seconds of fixation cross, the dynamometer was removed by the experimenter and both hands rested on the lap. Once the data was processed, the two blocks of each condition (OKS, OKS+L, OKS+R) were compared to each other. Since no significant differences were found between the two sets of blocks, each set was averaged together for final analysis.

### 2.4 Physiological measurements

#### 2.4.1 EEG and EOG

EEG was amplified using a BrainAmp DC amplifier (Brain Products, Munich, Germany) with a sampling rate of 1000 Hz, a band pass filter from 0.1 to 250 Hz and a notch filter at 50 Hz. The data was collected with 64 sintered Ag/AgCl ring electrodes mounted on an elastic cap (EASYCAP GmbH, Germany). 61 electrodes of the international 10/10 system were used for EEG recording (FP1, FPz, FP2, AF7, AF3, AF4, AF8, F7, F5, F3, F1, Fz, F2, F4, F6, F8, FT7, FC5, FC3, FC1, FCz, FC2, FC4, FC6, FT8, T7, C5, C3, C1, Cz, C2, C4, C6, T8, TP7, CP5, CP3, CP1, CPz, CP2, CP4, CP6, TP8, TP10, P7, P5, P3, P1, Pz, P2, P4, P6, P8, PO7, PO3, POz, PO4, PO8, O1, Oz and O2). Two electrodes were used to record horizontal EOG, placed on the outer canthi of the left and right eyes. Vertical EOG was recorded from one electrode below the observer’s left eye and from FP1. All electrode impedances were kept below 5 kΩ, referenced online to the left mastoid (TP9), with the ground electrode on position AFz.

Subsequent EEG and EOG processing was carried out with Brain Vision Analyzer 2.1 (Brainproducts, Germany). Data was digitally filtered between 0.1 and 40 Hz with a notch filter enabled at 50 Hz using an infinite-impulse-response filter (Widmann et al., 2015). Visual inspection of the data allowed rejecting segments with large mechanical artefacts. Nystagmus and other eye movements’ artefacts in the EEG were corrected using an infomax ICA as implemented in Brain Vision Analyzer 2.1 (Bell and Sejnowski, 1995, Makeig et al., 1995), which ran on the entire dataset. High-energy components reflecting nystagmus, blinks or other eye movements were manually removed. Afterwards, data was re-sampled to a power of 2 (1024 Hz). Before segmentation, EEG data was converted to a reference-free scheme through a Surface Laplacian estimation, using a spline order *m*= 4 and smoothing constant λ = 0.00001. Such parameters serve to reduce volume conduction, attenuating the effect of any remaining nystagmus artefact on the EEG and enhancing the location precision of experimental alpha modulations.

The data was segmented into each condition: BLC, BLO, OKN, OKN+L and OKN+R, each segment lasting 45 sec. Time-frequency analysis was implemented on pre-selected electrodes C3-C4 for the motor system, and O1-O2 for the visual system. The segmented blocks were convolved with 10-cycle complex Morlet wavelets, scaled logarithmically in 30 steps between 5 and 15 Hz. These parameters were set to maximize frequency precision over time precision. For statistical analysis, the spectral amplitude of 6 wavelet layers corresponding to the upper-alpha band were extracted, with central frequencies of 10.27, 10.67, 11.08, 11.51, 11.95 and 12.41 Hz, across the whole 45 seconds of the corresponding block. The 6 layers were then averaged together for each subject. The resulting average at each electrode was then pooled together for the motor region (C3-C4) and visual region (O1-O2).

To achieve a topographic representation of our results, scalp-maps were derived from fast-fourier transformations (FFT) over the entire 45 second segments, and contrasted between experimental conditions. To compute FFT, each 45 sec segment was segregated into 2 sec contiguous epochs, spectral amplitudes were extracted using a Hamming window with 50% overlap between contiguous epochs, allowing a spectral resolution of 0.5 Hz. All epochs within each of the 45 segments were averaged together at each electrode to create the final spectral distribution. Since FFT misses the temporal dynamics of cortical modulations along 45 seconds, no statistical analyses were performed on FFT data, only on the wavelet transformed data.

#### 2.4.2 EMG and Grip Force

Surface EMG was recorded using an ExG bipolar amplifier (Brain Products, Germany) connected to the EEG amplifier, sharing recording settings. Ag/AgCl electrodes were placed in bipolar montage on the skin over the flexor digitorum superficialis (FDS) of the left and right arm, with the ground placed near the lateral epicondyle of the elbow of the corresponding arm. Handgrip force was assessed with a digital hand dynamometer (Vernier, Beaverton, USA) connected to an independent computer running the Vernier Graphical Analysis software. The sampling rate was 10 Hz. One participant’s grip force data and another participant’s EMG data were lost through technical error. For analysis the EMG data of the maximum voluntary motor contractions (MVC) and the 45 sec periods of OKS-dynamic-handgrip of each hand were rectified and exported. MVC for both EMG and dynamometer grip force recordings was defined as the maximum peak achieved during the maximal contraction period of the respective hand. For dynamic handgrip, all data points of the 45 sec block were averaged together and normalized to the corresponding left or right MVC.

#### 2.4.3 Eye tracking

Eye movements were recorded by head-mounted video-oculography of the left eye with a sampling rate of 220 Hz (EyeSeeCam system, EyeSeeTec, Germany). The data was analysed offline through a custom MATLAB script, where blinks were first removed before detection of nystagmus quick phases, defined as eye velocity greater than 10 °/s and absolute acceleration greater than 300°/s^2^. The start point was determined at the time of peak velocity. The end point was considered when the eye velocity values reached 0°/s, indicating a change in the eye direction and the beginning of the slow phase (SP) of nystagmus used for analysis.

### 2.5 Data analyses

We first analysed the effects of simultaneous OKS-dynamic handgrip, where upper-alpha amplitudes were submitted to a 2×3 repeated measures ANOVA with factors “region” (motor, visual) and “condition” (OKS,OKS+L,OKS+R). The Greenhouse-Geisser correction was used when necessary to adjust for violations of sphericity. Afterwards, the relevant hypotheses stated in Table 1 were tested through directed post-hoc *t*-tests, with effect-sizes assessed through Cohen’s dz (Lakens, 2013). For the slow-phase velocity of OKN, a repeated measures ANOVA with the factor “condition” (OKS, OKS+L, OKS+R) was implemented with subsequent post-hoc *t-*test comparisons pertinent to experimental hypotheses. To control for the amount of muscular activation, the normalized grip-force and EMG levels were compared during OKS+L and OKS+R, expecting no significant differences between them.

Secondly, we analysed the effects of increasing visual input and OKS alone by submitting the upper-alpha amplitudes to a 2×3 ANOVA with repeated measures on the factors “region” (motor,visual) and “condition” (BLC, BLO, OKS). Likewise, the stated hypotheses in Table 1 regarding increasing visual input and OKS alone were tested through directed pairwise *t*-tests.

## 3. Results

Fig. 1 displays the wavelet-derived upper-alpha amplitudes at each analysed electrode grouped for motor and visual regions, along all experimental conditions. Each condition adds incremental load on the participant from resting with eyes closed in BLC, opening the eyes in BLO, then perceiving OKS, and finally simultaneous OKS and dynamic handgrip with left in OKS+L or right in OKS+R. Fig. 2 presents the contrasts between these conditions accompanied by a topographic representation derived from FFT.

**Fig. 1.**
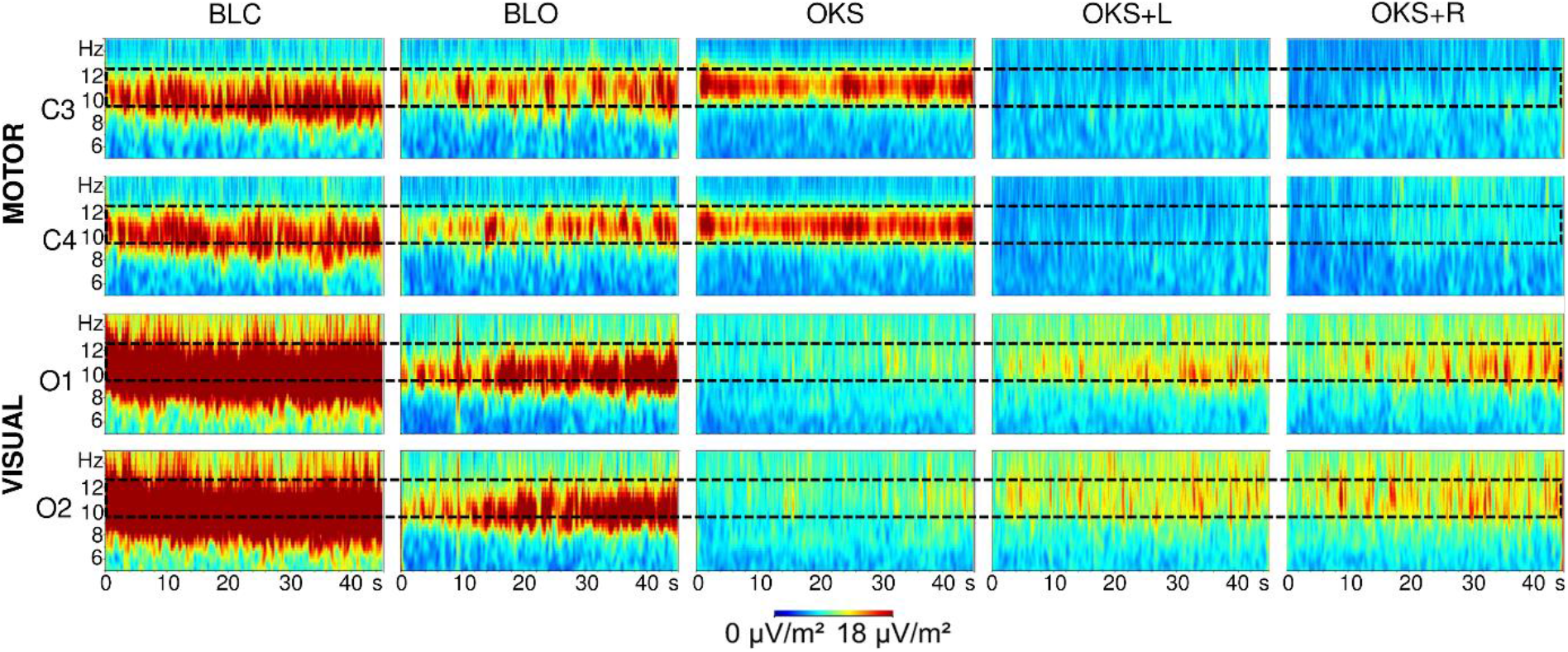
Grand average alpha amplitudes at the motor electrodes C3-C4 (motor region) and visual electrodes O1-O2 (visual region) extracted through 45 sec wavelet analysis of the different experimental conditions. The dashed rectangle highlights the upper-alpha sub-region (10-12 Hz) used for statistical analysis. The conditions are presented from less, to more load on the participant: BLC = Baseline closed eyes; BLO = Baseline open eyes; OKS = Optokinetic stimulation; OKS+L = Optokinetic stimulation + Left handgrip; OKS+R = Optokinetic stimulation + Right handgrip.

**Fig. 2.**
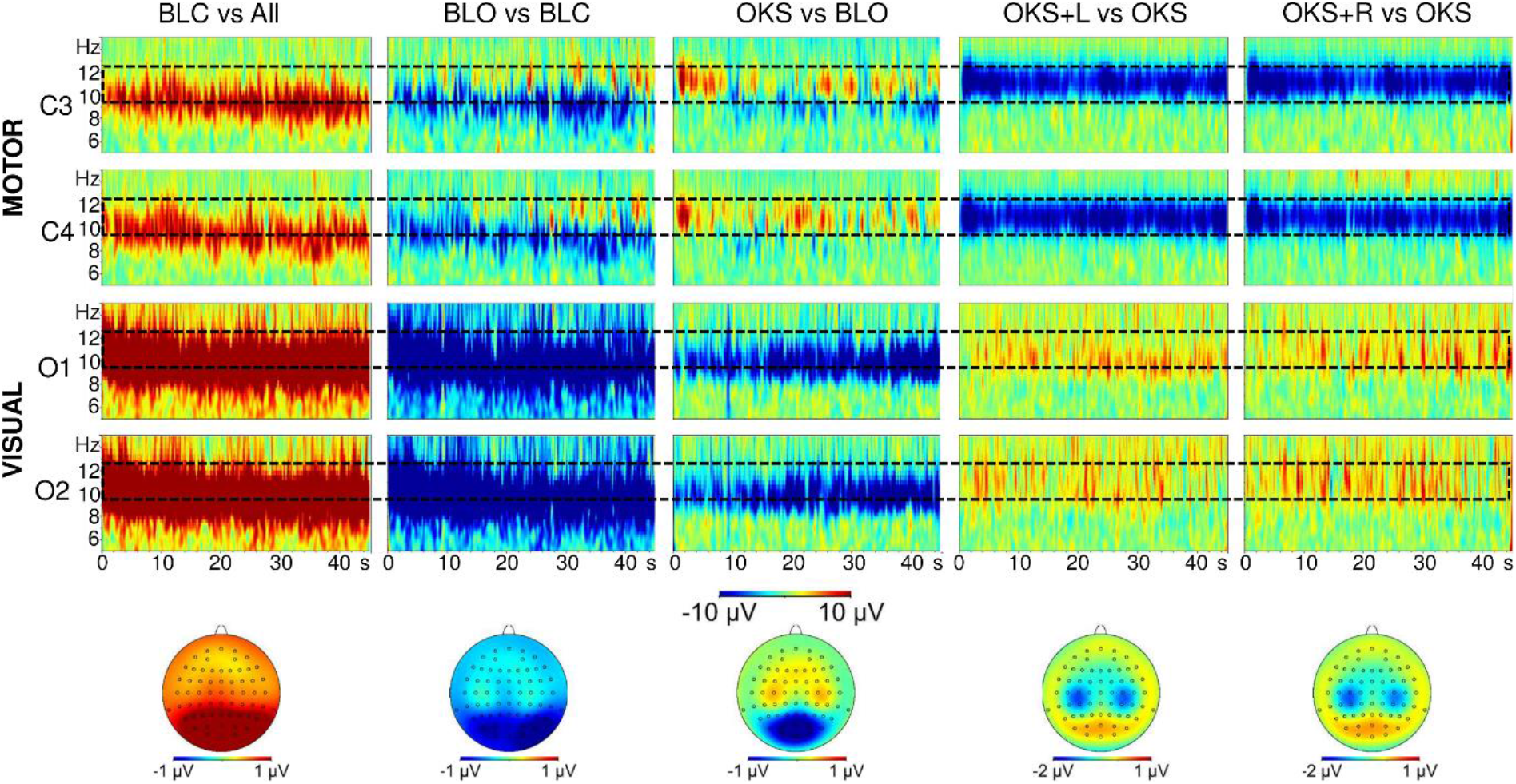
Contrasts of grand average alpha amplitude levels between consecutive experimental conditions. The conditions are presented from less, to more load on the participant: BLC vs all = Baseline closed eyes minus the average of all other conditions. BLO vs BLC = Baseline eyes open minus baseline eyes closed; OKS vs BLO = optokinetic stimulation minus baseline eyes open. OKS+L vs OKS = Optokinetic stimulation with left dynamic handgrip minus optokinetic stimulation. OKS+R vs OKS =Optokinetic stimulation with right dynamic handgrip minus optokinetic stimulation. Time-frequency representations are derived from wavelet analysis, grouped as in Fig. 1. The topographic representations below are derived from FFT over the entire blocks.

### 3.1 Effects during simultaneous OKS-Dynamic Handgrip

First the effects of handgrip were analysed to assess reliable EEG measurement. A significant effect of “condition” was observed [F(1.041, 13.53) = 5.92, p =.0285, η^2^ =.313], showing that OKS, OKS+L and OKS+R produced differences in upper-alpha amplitude. Table 2 shows the means and standard deviations of upper-alpha amplitude. Directed post-hoc t-tests are summarized in Table 3. They show that OKS+L and OKS+R significantly reduced upper-alpha amplitude at motor regions compared to OKS. This implies that dynamic-handgrip with either hand produced a detectable activation of motor areas, confirming our first hypothesis that motor activation is detectable during nystagmus, and supporting the reliability of EEG assessment in this condition. This amplitude reduction can be clearly appreciated in Fig. 1, where the motor alpha oscillations in OKS disappear during OKS+L and OKS+R. The contrast in Fig. 2 further reflects this dramatic reduction in OKS+L vs OKS and OKS+R vs OKS. As seen on Table 3, *t*-test comparisons for the visual region showed significantly greater alpha amplitudes during OKS+L and OKS+R than during the OKS condition. This confirms our second hypothesis, where handgrip execution had an inhibitory influence over the visual cortex even during visual stimulation. In Fig. 1, it can be appreciated how the introduction of OKS makes alpha disappear at the occipital locations O1 and O2, but return during OKS+L and OKS+R. The contrast in Fig. 2 further emphasizes greater alpha amplitudes at O1 and O2 in OKS+L vs OKS and OKS+R vs OKS.

**Table 2.** Mean (SE) upper-alpha amplitudes for each electrode averaged for 45 seconds at each condition. OKS = Optokinetic stimulation; OKS+L = OKS + Left handgrip; OKS+R = OKS + Right handgrip.

|  | **OKS** | **OKS+L** | **OKS+R** |
| --- | --- | --- | --- |
| **Motor region** | 14.00 (2.73) | 5.74 (.67) | 6.09 (.64) |
| **Visual region** | 8.19 (1.09) | 10.67 (1.44) | 11.06 (1.59) |

**Table 3.** *t*-scores and effect sizes for differences in upper-alpha amplitude between the isolated OKS with each of the OKS+Left and OKS+Right handgrip.

|  | **OKS vs OKS+Left** |  | **OKS vs OKS+Right** |  |
| --- | --- | --- | --- | --- |
| | $t(13)$ | $d_z$ | $t(13)$ | $d_z$ |
| **Motor region** | 3.37** | .90 | 3.29** | .88 |
| **Visual region** | -3.70** | .99 | -3.90** | 1.04 |

\* indicates significance $p < .05$
\*\* indicates significance $p < .01$

An interaction between “condition” and “region” [F(1.031, 13.41) = 14.60, p =.0019, η^2^ =.529], indicates that the experimental conditions also induced amplitude differences between the regions. This was driven by the effects described above having opposite directions, where the motor electrodes C3-C4 decreased alpha amplitude during simultaneous OKS and handgrip, and visual electrodes O1-O2 increased it. No main effect was observed for “region” [F(1, 13) = 1.74, p =.2103, η^2^ =.118] indicating no mean difference in amplitudes between regions across all experimental conditions.

Regarding OKN, a significant effect for “condition” was observed [F(1.40, 18.18) = 4.20, p <.0438, η^2^ =.244], indicating differences in the slow-phase velocity of OKN between the experimental conditions. Post-hoc *t-*tests indicated a significant increase during OKS+Left (M=15.44, SE=1.42) compared to OKS (M=13.40, SE=1.46), *t*(13)=-2.75, p=.0165, *dz =*-0.74. This is contrary to our third hypothesis, which predicted that the alpha enhancement produced by handgrip would be related to a *reduction* in slow-phase velocity. The effect was specific to left handgrips, as OKS+Right (M=14.65, SE=1.40) was not significantly faster than isolated OKS *t*(13)= −1.43, p=.1777, *dz =* −0.38. Meanwhile, the slow-phase velocity did not differ between OKS+Left and OKS + Right, *t*(13)= 1.78, p=.0973, *dz =* 0.48.

Regarding gripping force, there were no significant differences between %MVC for the left (M=27.95, SE=2.41) and for the right hand (M=24.88, SE=1.40), *t*(12)=1.51, p=.1572, *dz = 0.42.* In EMG, there was also no significant difference %MVC for the left (M=17.31, SE=2.79) than for the right hand (M=21.29, SE=2.86), *t*(12)= −1.79, p=.0995, *dz =* −0.50. In this way, gripping force and EMG measures suggest that participants exerted comparable effort with each hand, ruling out any confounding difference in force.

### 3.2 Effects during increasing visual input and OKS alone

Having confirmed that the EEG assessment during OKN is reliable, we proceeded to analyse the effects of increasing visual input and OKS alone over visual and motor regions. Repeated measures ANOVA over upper-alpha amplitudes revealed a significant effect of “condition” [F(1.18, 15.37) = 41.01, p <.0001, η^2^ =.76]. As seen in Table 4 and Table 5, post-hoc *t*-tests revealed significantly lower amplitudes in the visual region on BLO than BLC, and OKS than BLO. This confirms the fourth hypothesis that, besides alpha reduction after eye opening, OKS would produce further alpha reduction, reflecting visual activation as in previous EEG and fMRI studies. Fig. 1 shows how elevated alpha levels at visual electrodes O1 and O2 during BLC are reduced during BLO and disappear during OKS. Fig. 2 contrasts confirm these observations.

**Table 4.** Mean (SE) upper-alpha amplitudes for each electrode averaged for 45 seconds at the conditions: BLC= Baseline eyes closed BLO = Baseline open eyes; OKS = Optokinetic stimulation.

|  | **BLC** | **BLO** | **OKS** |
| --- | --- | --- | --- |
| **Motor region** | 13.38 (2.32) | 12.32 (2.48) | 14.00 (2.73) |
| **Visual region** | 30.43 (4.34) | 14.26 (2.01) | 8.19 (1.09) |

**Table 5.**
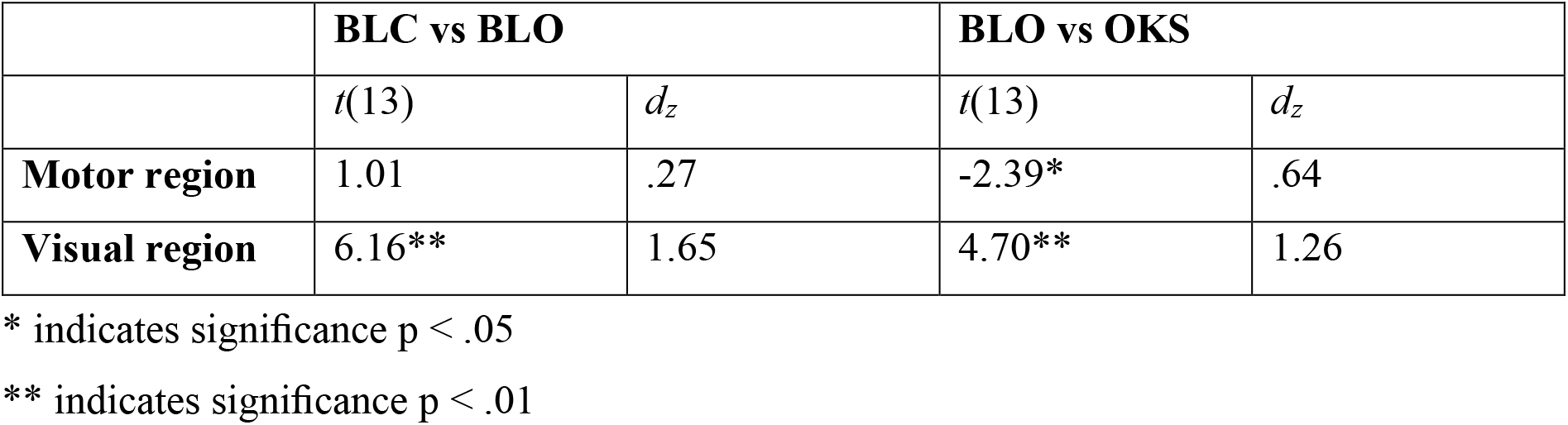
*t*-scores and effect sizes for differences in upper-alpha amplitude between the conditions with increasing visual input. BLC = Baseline closed eyes; BLO = Baseline open eyes; OKS = Optokinetic stimulation.

|  | **BLC vs BLO** |  | **BLO vs OKS** |  |
| --- | --- | --- | --- | --- |
|  | *t*(13) | *d<sub>z</sub>* | *t*(13) | *d<sub>z</sub>* |
| **Motor region** | 1.01 | .27 | -2.39* | .64 |
| **Visual region** | 6.16** | 1.65 | 4.70** | 1.26 |

\* indicates significance $p < .05$
\*\* indicates significance $p < .01$

A comparison between the BLC and the BLO conditions at the motor region showed no significant amplitude enhancement (Table 5), so that inhibition over the motor area was not evident after eye opening. However, when comparing BLO to OKS, a significant enhancement did appear, so that visual stimulation enhanced motor upper-alpha amplitude. In this way, our fifth hypothesis, that increasing visual input would produce greater inhibition over motor areas was only partially confirmed. However, a close inspection of Fig. 1 suggests that increasing visual input by eye opening in BLO and further OKS lead alpha oscillations to concentrate specifically in the inhibitory upper-alpha range, although perhaps BLO amplitude did not always surpass the broad alpha levels during BLC. When averaging the amplitude of each frequency layer in Fig. 1 across the entire 45 sec of each experimental condition (Fig. 3), it can be confirmed that BLO and OKS peak at faster frequencies than BLC, despite BLO not surpassing BLC amplitude on average. Such peaks also lie within the inhibitory upper-alpha region. These observations prompted us to analyse changes in peak frequency (rather than amplitude) over motor electrodes with increasing visual input.

**Fig. 3.**
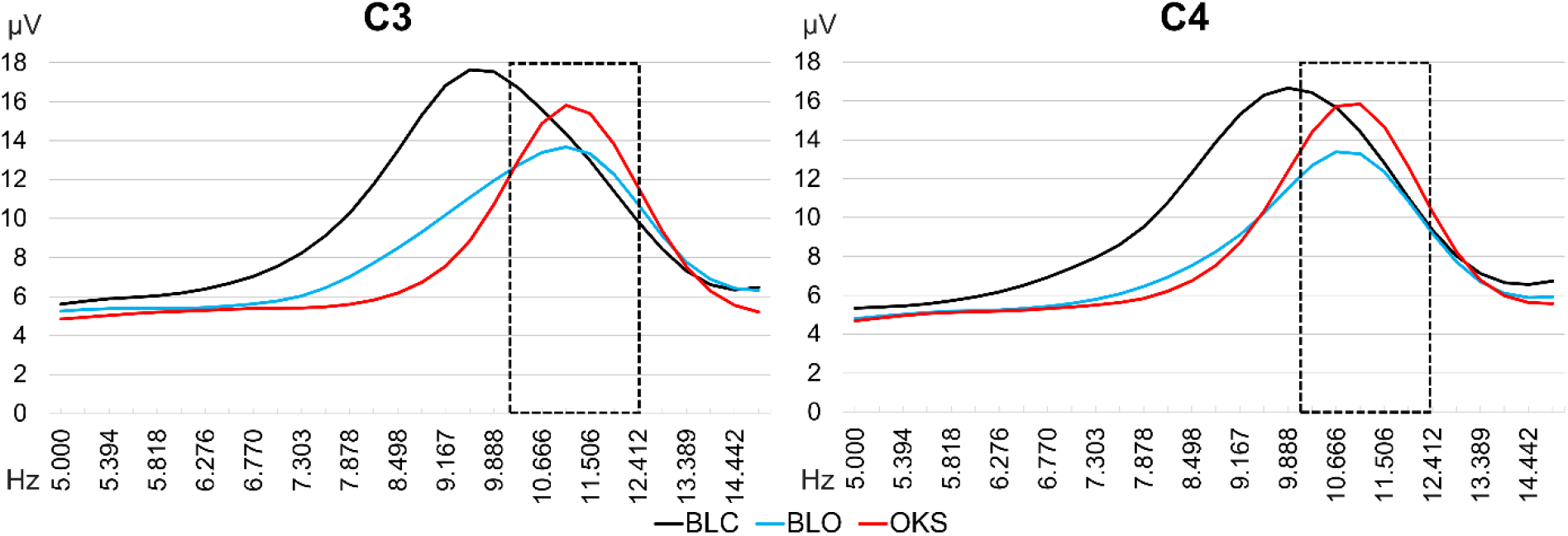
Amplitude distribution extracted from the grand average of wavelet analysis at motor electrodes C3 and C4. All time points along the 45 sec of each condition were averaged together. The dashed rectangle highlights the upper-alpha regions used for analysis. BLC = Baseline closed eyes; BLO = Baseline open eyes; OKS = Optokinetic stimulation.

### 3.3 Peak frequency changes over motor regions with increasing visual input

Fig. 4 displays the peak frequency at every data point (Wei, 2007) from the grand averaged time-frequency analysis displayed in Fig. 1 for the BLC, BLO and OKS conditions over motor electrodes C3 and C4. It can be observed that slower peak frequencies of around 8 to 10 Hz during BLC, broaden their range to include faster frequencies during BLO and finally concentrate around 11 Hz during OKS. After extracting the individual peak frequencies from each participant’s wavelet analysis at each condition, we implemented a 3×2 repeated measures ANOVA with the factors “condition” (BLC, BLO, OKS) and “electrode” (C3 and C4). Results showed a main effect of “condition” [F(2, 26) = 12.47, p =.0002, η^2^=.49], implying that each of the experimental conditions produced a change in the peak frequencies at each electrode. Table 6 displays the mean peak frequencies at each phase, while Table 7 shows the results of post-hoc *t*-tests. It can be observed that BLO and OKS showed significantly faster peak frequencies than BLC, within the upper alpha range. Meanwhile, peak frequencies did not differ between BLO and OKS. These observations support that increasing visual input biases oscillations towards the inhibitory upper-alpha range. As seen in the previous section, once in the upper-alpha range, greater visual input increases upper-alpha amplitude.

**Fig. 4.**
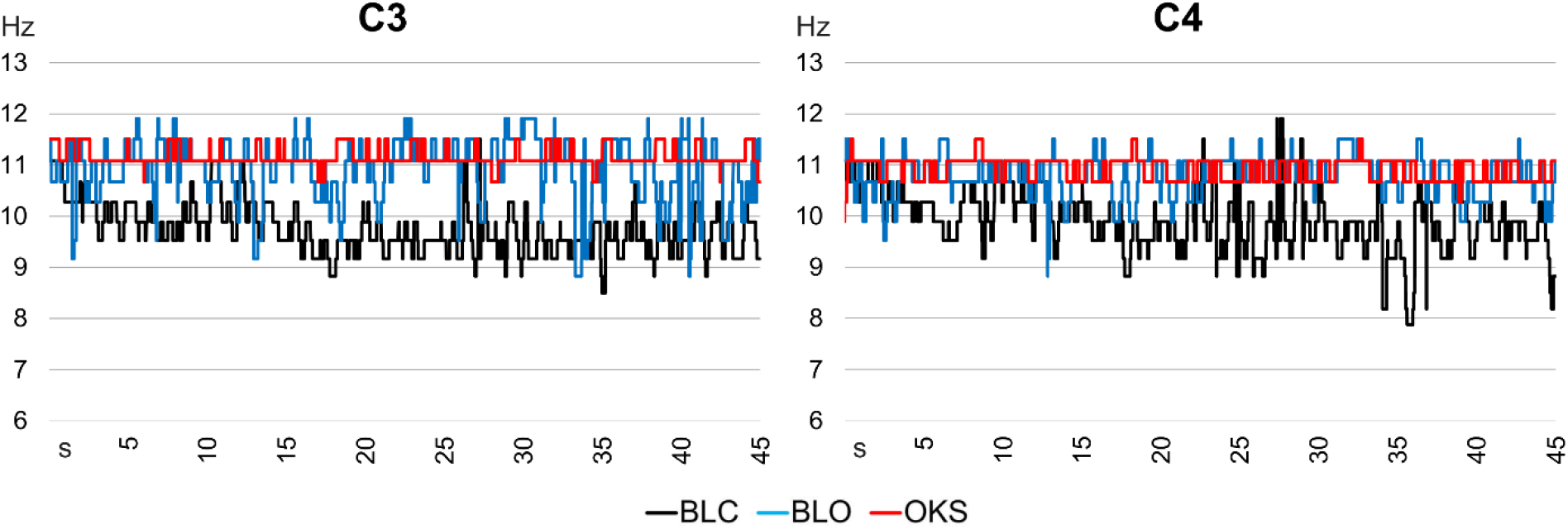
Peak frequencies at every time point of the grand-average wavelet analysis in the motor electrodes C3 and C4 during the experimental conditions with increasing visual input. The conditions are overlaid. BLC = Baseline closed eyes; BLO = Baseline open eyes; OKS = Optokinetic stimulation.

**Table 6.** Mean (SE) peak frequencies for each electrode averaged for 45 seconds at the conditions: BLC= Baseline eyes closed BLO = Baseline open eyes; OKS = Optokinetic stimulation.

|  | **BLC** | **BLO** | **OKS** |
| --- | --- | --- | --- |
| **C3** | 10.55 (.26) | 11.44 (.23) | 11.38 (.16) |
| **C4** | 10.62 (.22) | 11.26 (.19) | 11.20 (.18) |

**Table 7.** *t*-scores and effect sizes for differences in peak frequency between the conditions with increasing visual input. BLC = Baseline closed eyes; BLO = Baseline open eyes; OKS = Optokinetic stimulation.

|  | **BLC vs BLO** |  | **BLC vs OKS** |  | **BLO vs OKS** |  |
| --- | --- | --- | --- | --- | --- | --- |
| | $t(13)$ | $d_z$ | $t(13)$ | $d_z$ | $t(13)$ | $d_z$ |
| **C3** | -4.45** | 1.19 | -4.23** | 1.13 | 0.36 | 0.10 |
| **C4** | -3.31** | 0.88 | -2.92* | 0.78 | 0.57 | 0.15 |

\* indicates significance $p < .05$
\*\* indicates significance $p < .01$

An interaction between “electrode” and “condition”[F(2, 26) = 3.44, p =.0474, η^2^=.21] indicates that the experimental conditions introduced differences in peak frequencies between the two electrodes, but post-hoc analyses showed that these differences were not practically meaningful (e.g. C4 during BLO greater than C3 during BLC). Meanwhile, peak frequencies at each electrode did not differ from each other during BLC (*t*= -.85; *p*=.4109), BLO (*t*= 1.30; *p*=.2173), or OKS (*t*= 1.41; *p*=.1804), indicating that the modulations in peak frequency occurred in parallel at both electrodes.

## 4. Discussion

Optokinetic stimulation and nystagmus reduced EEG upper-alpha band amplitude over occipital electrodes, reflecting activation, and also enhanced upper-alpha over central motor electrodes, reflecting inhibition. Similarly, simply having the eyes open produced a significant shift in motor peak frequency towards upper-alpha (compared to eyes closed), which then increased amplitude during OKS. This suggests that increasing amounts of visual input lead to increasing levels of motor inhibition. When participants performed dynamic handgrip during OKS, the opposite pattern emerged: bilateral motor alpha reduction, and an occipital increase. This implies that generating motor activity during OKS disrupted the occipital activation normally induced by visual stimuli. Behaviourally, OKN slow phase velocity increased only during left handgrip, despite no difference in gripping force between hands. Together, these results support that upper alpha band oscillations reflect the inhibitory interactions between visual and hand/motor systems, affecting oculomotor responses. Additionally, this extends previous EEG studies by showing that these inhibitory interactions hold when both systems are simultaneously and continuously engaged.

### 4.1 EEG reliability during nystagmus

A necessary first point to address regards the reliability of EEG assessment during nystagmus. To validate this, we tested whether dynamic handgrip during OKS could reproduce the robust bilateral motor alpha amplitude reduction (cortical activation) classically found during handgrip and other hand movements (Cross-Villasana et al., 2016, Deiber et al., 2001, Gerloff et al., 1998, Pfurtscheller and Lopes da Silva, 1999). When comparing handgrip-OKS against OKS alone, this observation was indeed replicated (Fig 2). This shows that any artefact produced by nystagmus over the EEG need not obscure the alpha-band modulations induced by handgrip. The EEG was refined with ICA to subtract the nystagmus artefact (Bell and Sejnowski, 1995, Makeig et al., 1995), and Surface Laplacian to reduce volume conduction (Kayser and Tenke, 2015, Tenke and Kayser, 2005). In this way, it was also possible to demonstrate the activation of the visual cortex during OKS, (reflected by an alpha-amplitude decrease), as in previous OKS-EEG (Gulyás et al., 2007, Magnusson et al., 1985) and fMRI studies (Becker-Bense et al., 2006, Bense et al., 2001, Dieterich et al., 2003, Kikuchi et al., 2009). With the reliability of EEG confirmed, we proceeded to analyse the interactions between the visual and motor systems, their effect on nystagmus, and motor inhibitory effects produced by OKS.

### 4.2 Motor inhibitory influence over the visual cortex

During simultaneous handgrip-OKS, apart from motor activation, we detected the increase in occipital alpha synchrony (Fig. 2) previously found during isolated hand movements (Cross-Villasana et al., 2016, Deiber et al., 2001, Pfurtscheller and Lopes da Silva, 1999). This implies that handgrip inhibits visual cortex, despite continuous stimulation by the optokinetic visual display. This is supported by the suggested inhibitory function of alpha band oscillations (Bazanova and Vernon, 2014, Jensen and Mazaheri, 2010, Klimesch et al., 2007, Pineda, 2005, Romei et al., 2008, Sauseng et al., 2009), especially in the upper-alpha sub-band (Hummel et al., 2002, Klimesch et al., 2007, Pfurtscheller and Lopes da Silva, 1999, Zarkowski et al., 2006). While the posterior alpha enhancement during simultaneous OKS and dynamic handgrip resembles that of isolated dynamic handgrip (Cross-Villasana et al., 2016), in the context of OKS it implies that visual cortical activation was interfered with by the repetitive handgrip movement. For that reason, this observation complements previous electrophysiological evidence where one active thalamocortical stream can be interrupted by the activation of a second stream (Crabtree and Isaac, 2002, Zikopoulos and Barbas, 2007), here the visual stream being interrupted by the motor stream. This suggests that these inhibitory interactions (potentially mediated initially at the thalamus) can be observed in the scalp EEG as an increase in alpha power (Hughes and Crunelli, 2005, Larson et al., 1998, Vijayan and Kopell, 2012) reflecting the interfered thalamocortical stream (Pfurtscheller and Lopes da Silva, 1999, Steriade, 2000, Sterman, 1996).

Converging evidence supports the idea that limb motor activation can interfere with visual processing. For example, the latency of saccade execution towards a target is slower whenever it is accompanied by a hand movement, suggesting that the hand movement interfered with the visual processes that lead to the saccade (Sailer et al., 2015). Research in cat and monkey has shown that periods of increased visual attention are accompanied by motor inhibition, muscle tone reduction and motor cortical down-regulation, presumably to reduce motor interference (Rougeul et al., 1979, Sterman, 1996). In humans, O’Muircheartaigh et al. (2015) observed in a large diffusion-weighted and resting-state fMRI sample that visual and motor thalamocortical streams are strongly anti-correlated. When Kober et al. (2015) trained participants to enhance the EEG sensorimotor-rhythm (12-15 Hz) by reducing their resting muscular activation, they observed improved performance and enhanced event-related potentials in a visual short-term memory task. After the training, Kober et al. also observed reduced resting EEG functional connectivity between motor and parieto-occipital areas. Together, those findings suggest that reduced interference between motor and visual areas lead to improved visual stimulus processing and task performance (Kober et al., 2015). In contrast, we here produced a deliberate interference of the visual system by the motor system.

### 4.3 Effect of dynamic handgrip on OKN

Together with visual alpha enhancement, dynamic-handgrip increased OKN slow phase velocity, and furthermore only with the left hand. Given that nystagmus was sped up despite the disruption of the visual cortex indicated by alpha synchrony, this implies that limb motor activation may affect oculomotor processes independently of its effects on the visual system. In the light of recent work, we suggest this occurs via an indirect route that stimulates the oculomotor network generating the nystagmus. One candidate would be the dorsal attentional network, key parts of which are activated during OKN, such as the frontal eye fields (FEF; Ruehl et al., 2019). In a recent TMS study, disrupting the FEF during OKN slowed the eye movement (Mastropasqua et al., 2020). Here we can apply the same logic in the opposite direction: if intensive hand gripping facilitated the dorsal attentional network, this could account for the speeding of slow phase velocity of OKN. Hand movements have shown to influence frontal regions beyond the motor cortices (e.g. Rektor et al., 2006; Derambure et al., 1997). Future research that investigates the time course of modulations produced by single handgrips across the scalp during OKS may provide more insight into these questions.

Although the effect of handgrip on OKN was apparent for the left hand and not the right hand, there was no statistical difference between left and right hand effects in a direct comparison, meaning future work is necessary to determine how hand-specific this effect was. Given that our participants were right-handed, but that there was no significant difference between grip force applied between left and right hands (confirmed here with a dynamometer), this could imply that greater neural modulations were involved during left hand movements (Cross-Villasana et al. 2016), possibly due to greater effort being required from right handers, leading to greater indirect effects (e.g. via the attentional system) on OKN. Relatedly one could postulate that the observed pattern may arise if there is a putative right hemispheric bias of higher vestibular function in right handers (Dieterich and Brandt, 2018). If inter-hemispheric competition controls vestibular cortical processing (Arshad, 2017, Bloom and Hynd, 2005), left hand movements mostly controlled by the right hemisphere may strenghten the right hemisphere amid the competition. Such an account, however, also requires future work to demonstrate that left handgrip leads to stronger inter-hemispheric competition between vestibular processes.

### 4.4 Inhibitory influence over the motor cortex with increasing visual input

Eye opening sped up the peak frequency of the EEG oscillation over the motor cortex, which was on average 1 Hz faster than in the eyes closed condition, putting it within the inhibitory upper-alpha range. The introduction of OKS further increased the amplitude of this new peak, in addition to eliciting the expected visual activation (Fig 3; Fig 4). This suggests that OKS lead to the simultaneous inhibition of the motor system, and activation of the visual system, in accordance with previous OKS-fMRI observations (Becker-Bense et al., 2006; Ruehl et al. 2019), and with EEG studies using other kinds of visual stimuli (Brechet and Lecasble, 1965, Koshino and Niedermeyer, 1975, Pfurtscheller and Lopes da Silva, 1999). Furthermore, considering that the upper-alpha band is particularly associated with inhibition (Hummel et al., 2002, Klimesch et al., 2007, Pfurtscheller and Lopes da Silva, 1999, Zarkowski et al., 2006), it can be proposed that increasing visual input, from closed eyes, to eye opening, to visual stimulation, produces progressively greater inhibition of the motor cortex. This argument is supported by research using transcranial magnetic stimulation (TMS), where changes in the state of the visual cortex affect motor-cortical excitability. For example, a TMS pulse to the visual cortex reduces the motor-evoked-potential produced by a second pulse to the motor cortex (Strigaro et al., 2015). Furthermore it has been shown that simply opening the eyes reduces motor excitability, while putting on a blind-fold increases it (Leon-Sarmiento et al., 2005, Strigaro et al., 2015). Overall, these parallel findings in EEG, fMRI and TMS support that the current upper-alpha peak at eye opening, and its further enhancement during OKS, do represent a state of motor down-regulation produced by visual activity.

### 4.5 Implications

Whole scalp EEG holds great potential as a valuable tool in research fields involving nystagmus, such as vestibular research, since it can compensate for the lack of temporal resolution in other techniques currently used in the field (Lopez and Blanke, 2011). OKS and OKN are also used in other fields such as in binocular rivalry where it has been previously suggested that simultaneous EEG should be possible using ICA (Einhäuser et al. 2017). Our whole-scalp recording during OKS and use of ICA and Surface Laplacian supports the successful use of EEG when nystagmus artefacts are unavoidable. Eye-movement artefacts can be effectively dealt with these techniques, allowing new areas of EEG analysis (e.g. Cross-Villasana et al., 2018, Delorme et al., 2012, Delorme et al., 2007). The use of electrooculography to detect nystagmus artefacts on EEG, and a large density of electrodes for better spatial filtering is recommended to reliably isolate the signal of interest through these techniques.

A better understanding of sensorimotor interactions could help in the development of rehabilitation strategies that rely on sensory input and/or OKS for treating vestibular and attentional disorders (Kerkhoff et al., 2012; Cousins et al., 2014, Karnath and Dieterich, 2006). Down-regulating the visual cortex during OKS exposure through handgrip exercise could accelerate the desensitization to visual motion (Cousins et al., 2014, Guerraz et al., 2001), or help retrain reliance on the vestibular system for keeping balance (Cousins et al., 2014).

### 4.6 Limitations

The current experimental block-design was based on previous imaging studies of OKS, expanding upon their findings using EEG wavelet analysis. Our block design and wavelet parameters were designed prioritising frequency precision over temporal precision to better detect upper-alpha modulations. Future event-related studies relying on time-frequency analysis (but forfeiting frequency precision for greater time precision) will be necessary to investigate the time-course of the inhibitory effects produced over the visual cortex after each handgrip, or those over the motor cortex during OKS. A recent study reported that frontal eye field TMS during OKS modulated the anterior-posterior alpha EEG balance, the OKN slow phase velocity, and simultaneous perceptual processing, interestingly with no effects of TMS over primary motor cortex (Mastropasqua et al. 2020). It would also be important to determine which other cortical and subcortical areas are modulated after each handgrip, which may contribute to the currently observed acceleration of the slow-phase velocity of OKN.

## Abbreviations

OKS: Optokinetic stimulation
OKN: Optokinetic nystagmus

